# Diurnal variations in serum metabolites of wintering redheaded buntings

**DOI:** 10.1101/561407

**Authors:** Neelu Jain Gupta, Samya Das, Mrinal Das, Rakesh Arya, Ranjan Kumar Nanda

## Abstract

Daily behavioural and physiological changes in bird may reflect in biofluid metabolite composition. Locomotor activity, food intake and body temperature of group (n=7) of male migratory redheaded buntings held under short days (8L:16D, SD) were monitored besides blood sampling at midday (ZT4: 4 hours zeitgeber time starting ZT0 as lights ‘on’) and midnight (ZT16). The birds exhibited higher activity and increased feeding during daytime with negligible activity and feeding at night. Gas chromatography mass spectrometry and chemo-metric analyses of bird serum revealed higher levels of lipid (palmitic, oleic and linoleic acids) and protein (uric acid and proline) catabolites in daytime serum samples as compared to night samples. Higher night-time levels of short chain fatty acids indicated utilization of glucose and lipolysis in night fasted birds. High night-time levels of taurine, a sulphur amino acid has adaptive advantage to night migratory song birds. The diurnal differences in metabolite patterns suggests differential energy expenditure during day and renders survival benefit to buntings as night migrants. We propose a GCMS method that could be useful to unravel different annual life-history stages including migration.

## INTRODUCTION

Molecular basis of temporal regulation of metabolic homeostasis is extensively studied in different experimental animal models. Wild animals exhibiting seasonal behaviour can also be useful to delineate biochemical changes that distinguish seasonal metabolic phenotypes. Studies on migratory birds are important, because these birds, exhibit clear differences in their seasonal life history stages across a year (LHS such as wintering, breeding, migrating etc.; Malik et al., 2014); the behavioural and physiological changes during these annual LHS results from dynamic interplay of a photo-regulated endogenous circadian clock and functional metabolic states (FMS).

The transitions between LHSs in a seasonal species are accomplished by changes at multiple regulatory levels (Trivedi et al., 2014). These multi-level inputs from differential gene expression and post transcriptional and post translational modifications consummate into a thorough flux of information viz. metabolite fluctuations. A thorough understanding of FMS thus, requires comprehensive experimental analyses involving integrative analytical techniques, capable of identifying and quantifying metabolic compounds. Gas/liquid Mass spectrometry techniques (G/LC-MS) obtains comprehensive landscape of organismal metabolic homeostasis. A metabolomic assay is capable of efficiently summarizing functional metabolic states and an inverse assignment of metabolomics data to underlying processes can render insight into metabolic systems at various levels of molecular organization (Riekeberg & Powers, 2017). However, metabolomic approaches are limited by large number of metabolic compounds with differentially detectability in real-time studies. Deriving pertinent information from metabolomic dataset of migratory bird tissue sample is particularly complicated for two reasons-one, confidence of interpretation of particular metabolite in wild animal tissue and second, a longitudinal investigation on migratory bird correlating daily behavioural and physiology to downstream metabolic changes employing metabolomics tools requires comparative analyses of bird tissue from several LHS during an annual cycle, leading to enormous datasets to begin with. No doubt, molecular analyses of photo-regulation of migration pertain to functional genomics data and fewer studies have engaged metabolite assay of comparative FMS. A study showed that migratory fuelling in long distance migratory red knot (*Calidris canutus*) depended on daily and seasonal changes in three metabolites viz. triglycerides, uric acid and β-hydroxy-butyrate (Johnston et al., 2016).

To highlight the importance of metabolic profiling a GC-MS method needs to be standardized that could be useful to understand diurnal variation of metabolic dependence in night migratory songbirds i.e. redheaded bunting. Redheaded bunting (*Emberiza bruniceps*) is a night-migratory (breeding grounds 52°N) songbird, overwintering in India (Ali & Ripley, 1974; Jain & Kumar, 1995; Jain-Gupta & Kumar, 2013). Although, with a long-term goal to assess seasonal metabolic adaptation dynamics of songbirds, present report includes diurnal metabolic changes under short days; short days minimize daily differences in leptin, insulin, total serum proteins and urea (Soppela et al., 2008). To assess photo-regulated day and night metabolic changes in buntings during wintering LHS i.e. the birds were housed under short-days for 4 weeks and blood samples drawn at midday (ZT4) and midnight (ZT16). Daily activity, food eating pattern and body temperature were measured along 24 h to correlate with diurnal changes in serum metabolites. The wintering LHS represented a FMS with lower daily energy expenditure (Wikelski et al., 2008) and ruled out a scope for alterations in confounding factors such as body weight.

## MATERIAL AND METHODS

The animal care and procedures adopted in this study were as per guidelines of the Institutional Animal Ethics committee (IAEC) of Chaudhary Charan Singh University, Meerut. Adult male redheaded buntings (*Emberiza bruniceps*) were captured and brought to Ghaziabad (Ghaziabad, India: 28.6° N; 77.4° E) in February 2014 and allowed to acclimatize for 3 weeks in an outdoor aviary (size: length × breadth × width:2 m × 2 m × 1.2m). Birds were fed on cereal grains (*Setaria italica*) supplemented with gritted hard-boiled eggs and water was available ad libitum. A group of male birds (n=7) were transferred to short days (8L: 16D; SD) i.e. 8 hours of light: 16 hours of dark, light intensity 1.55 W/m^2^ during light phase and 0.002 W/m^2^ during dark phase, in individual cages (size: 45 cm × 30 cm × 30 cm) kept in well aerated photoperiodic chambers (size: 1 m × 1 m × 1 m). All birds had small sized testes (testis volume = 0.52–0.76 mm^3^) and were without any photo-stimulation as apparent from body mass (24-27 g) at the time of capture. Ambient temperature was monitored using Easy Log USB, Lascar electronics Inc. PA, USA and maintained constant at 22 °C. Birds that died within a week of experiment were excluded from analysis.

### Behavioural Assay

The locomotor activity of individually caged birds (n=7) was recorded using activity cages mounted with two perches and a top mounted infrared sensor system (LC- 120-PI make Digital Security Controls, Canada) detecting general activity of bird within the cage. The activity thus recorded by sensor in 5min bins was transmitted to data-logging system-Chronobiology Kit (Stanford Software System, USA) enabling channel-wise output as actograms (double plots of activity in successive days) and exported quantified bouts of activity/ minute for the duration of experiment.

### Physiological parameters and blood sampling for GC-MS analysis

Daily food eating pattern (DFEP), of buntings was observed for three days in continuum, wherein individually caged birds (n=7) were administered a fixed proportion of seed-egg mixture as food, by weighing and replenishing food bowls every 2 hours during lighted hours. Since the birds do not eat in dark hours, food was replenished only once during mid-night instead of two hourly measurements (Das & Jain-Gupta, 2016) Surface body temperature (SBT) was recorded 6 times in 24-hours using thermoscan (range 32-42.5 °C) at ∼1cm distance from the keel region skin surface exposed by gently blowing air. Birds were never handled manually except during 4 hourly SBT recording.

All birds were kept for 28 days in short days (8L: 16D) and after showing a synchronized photoperiodic properties, blood samples were collected at two time points representing midday (ZT4) and midnight (ZT16). The blood sampling sequence was randomized besides other care taken to avoid metabolic alteration due to physical stress, maintaining at least 36h gap between two subsequent sample collections. Bird’s wing vein was punctured with a fine sterile needle and blood droplets (yielding 100- 250 μL volume) were gently collected into heparinized microhematocrit capillary tubes. Blood samples were kept at 25 °C for 0.5 hours, centrifuged at 2,000× g at 4 °C for 10 min. Supernatant were collected as serum and aliquoted in coded microcentrifuge tubes and stored at −80°C until further analysis. Minimum two freeze thaw cycles were allowed before use of these materials.

### Quality Control (QC) and Randomization

Following recommended protocols, an equal volume of all study serum samples was pooled to prepare a quality control sample (QC) (Sangster et al., 2006; Dunn et al., 2011). Coded samples were randomized using a web-based tool (www.randomizer.org) to process those for metabolite extraction and derivatization in batches followed by GC-MS data acquisition within 24 hours. To minimize operator biasness, we adopted a double blinding approach during serum sample collection and metabolic profiling study. Two independent researchers carried out sample collection and GC-MS data acquisition.

### Serum Processing and Derivatization

Serum volume is a limiting factor for such studies so we first standardized appropriate serum volume (5, 10, 20, 30, 40 and 50 μl) to be taken for molecule profiling study using QC sample. Five to six test serum samples in a batch were brought to room temperature by thawing in ice, before undertaking TMS (trimethylsilyl) derivatization following the method described by Das et al., with minor modification (Das et al., 2015). In brief, 50μl deep freeze serum sample thawed on ice and 10μl of freshly prepared isoniazid solution (1 mg/ml) was added as an internal standard. A volume of 800 μL ice-cold methanol was mixed with the sample then vortexed for 30 sec. The suspension was centrifuged at 15,000 × g for 10 min at 4 °C and supernatant was dried in a speed vac at 40 °C. Dried sample was treated with 2% methoxyamine HCl in pyridine (MOX) reagent at 60 °C for 2h followed by a silylation step with N,O-Bis(trimethylsilyl)trifluoroacetamide (BSTFA) at 60 °C for 1h. After derivatization, sample tube was centrifuged at 10,000 × g for 5 min and supernatant was transferred into a vial insert kept inside a 2 ml screw capped glass vial.

### GC-MS Data Acquisition

Using an automatic sampler (MPS, Gerstel Germany), 1μl of derivatized test serum sample was loaded using split less mode to an RTx-5 column (5% diphenyl, 95% dimethylpolysiloxane; 30m x 0.25mm x 0.25μm; Restek USA) in a GC-TOF-MS (Pegasus 4, Leco, USA) for separation. Helium was used as carrier gas at a constant flow rate of 1 ml/min. The inlet temperature was fixed at 300 °C during injection, temperature gradients of 80°C to 150 °C (ramp of 7°C/min), 150 °C to 270 °C (ramp of 10°C/min) with a hold time of 1 minute between two ramps. After reaching final temperature an isothermal run for 15 minutes was maintained. Electron ionization (EI) mode was fixed at 70 eV to scan ions of 35 to 600 m/z ranges. Maximum scan speed was 20 Hz with a 230 sec solvent delay. The ion source temperature was fixed at 240 °C and data acquisition voltage was 1600. Sample introduction to data acquisition parameters in GC-MS were controlled through ChromaTOF software (Leco, USA) and the total run time was 2340 sec per sample.

### Data pre-processing and peak alignment

Raw GC-MS data (.peg) files of all the study samples (n=14) were pre-processed using ChromaTOF. Alignment of all GC-MS data files was carried out using ‘Statistical Compare’ feature of ChromaTOF. For peak picking, peak width was set at 1 sec and signal/noise (S/N) threshold was 100. For tentative molecular feature identification mainlib (2,12,961 spectra) and replib (30,932 spectra) libraries from NIST (version 11.0) were used with a minimum similarity index of 750. Maximum retention time deference was set at 0.5 sec and for mass spectral match minimum spectral similarity among aligned molecules was set at 600. Unsilylated molecules were removed from the data matrix manually. Aligned peak information was exported to .csv format and molecules absent in more than 50% of samples of at least one class (time point) were excluded from analysis. For metabolite analysis, both uni- and multi-variate analyses of the total area normalized metadata were carried out using MetaboAnalyst 3.0 to identify important deregulated molecules (Xia & Wishart, 2015). Missing values were imputed with half of the minimum value of study population and data matrix was normalized with peak area of the feature representing the internal standard isoniazid. Following which, generalized log transformation and universal scaling method were employed to obtain a near Gaussian distribution of the variables to carry out multivariate analyses (principal component analysis: PCA, and random forest: RF). Uni-variate analysis in terms of paired t-test (p<0.05) was carried out using MetaboAnalyst. Features qualifying criteria of t-test p value <0.05 and Mean Decrease Accuracy score > 0.002 of a Random Forest (RF) model were selected as important deregulated molecules. Selected important molecular list was taken for hierarchical clustering using Ward linkage and Euclidean cluster method and separately Principal Component Analysis (PCA) was carried out to see the clustering pattern.

### Confirmation of identity of important molecules

Identity of 47 molecules was confirmed using reported retention indices. Commercial standards of tentatively identified important molecules were oximated and silylated following similar methods used for analysing the samples. Derivatized standards were run following the GS-MS methods used for sample data acquisition. Retention time and fragmentation pattern of both commercial standards and important metabolites were matched to establish the identity of these important molecules.

### Pathway analysis

To identify deregulated molecular pathways, identified molecules were used as input list in MetaboAnalyst 3.0 (Xia & Wishart, 2015). Pathway library of *Gallus gallus* was used for pathway analysis.

### Statistical analysis

In behavioural study, student’s t-test was used to compare total activity and food intake between different observations. One-way ANOVA followed by post-hoc Newman-Keuls was used for comparing body temperature at different time points. Significance was taken at p<0.05. Data were plotted and statistical analyses performed using Prism GraphPad software (GraphPad ver. 5.0, San Diego, CA).

## RESULTS

### Circadian rhythm in behavioural parameters

Buntings were predominantly day-active (t_5_=15.52, p<0.0001) and activity onset and offset coincided with the lights-on and lights-off (Fig 1 A, B and C). There were two peaks in daytime activity and first peak hour in the morning was marked by almost double the activity of the second peak in the evening (Fig 1D) respectively. Buntings exhibited a bimodal pattern of food intake through the light phase; 2-hourly food consumption declined 11.4% from morning to afternoon followed by 19.1% increase from afternoon to evening (Fig 1 E and F). A daily rhythm was observed in surface body temperature (F_(5, 47)_=4.43, P=0.025, One-way ANOVA followed by Newman-Keuls Multiple post hoc Test) with an increase after lights-on (Fig 1 G and H). We observed significant circadian rhythm in activity, food intake and surface body temperature in buntings (Fig 1I).

**Figure 1:**
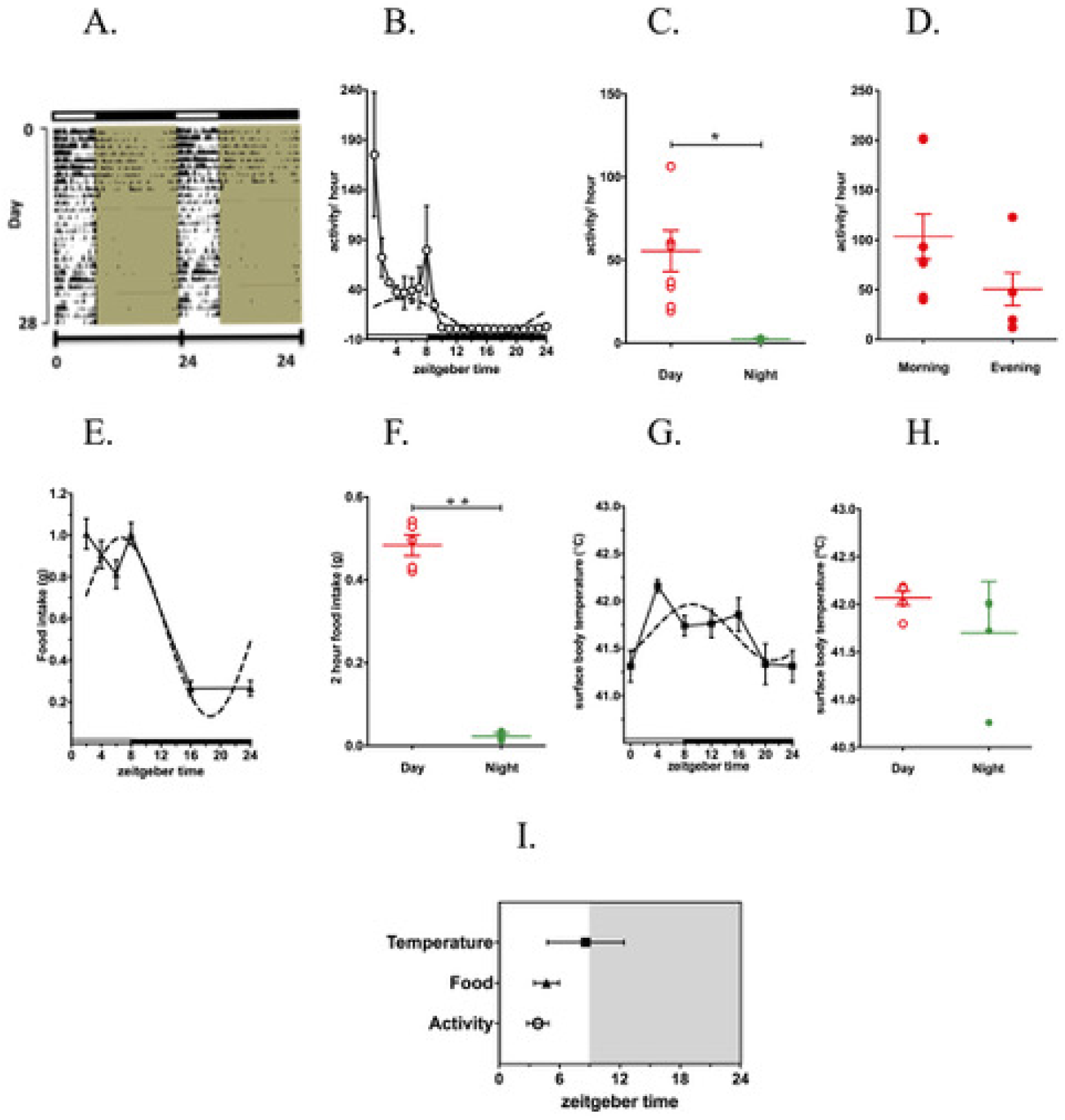
Diurnal variation in behaviour and physiology of male redheaded bunting*, Emberiza bruniceps* held under 8L: 16D. A. Double plotted actogram of a representative bunting. B. Daily activity profile of group (n=6). C. Comparison of hourly daytime and night-time activity. D. Comparison of morning-evening peaks of daytime activity. E. Daily food eating pattern DFEP of group (n=6). F. Comparison of hourly food intake during daytime (red) and night-time (green).G. Daily changes in surface body temperature. H. Comparison during daytime and night-time. I. The time of peak (acrophase) of activity rhythm, daily food intake rhythm and surface body temperature. Dotted lines represent the Cosinor regression curve drawn through the time points to deduce daily rhythms. The data for group are plotted mean ± SEM, p< 0.05.

### Serum volume standardization

Maximum identifiable molecular features using GC-MS during serum volume standardization were observed at 50 μl volume (S1). Identity of a sub set of these molecules (n=47) was confirmed using reported retention indices (S1 Table).

### Differences in chromatogram of serum samples collected during day and night

Fig 2 A and B show representative total ion chromatograms of bunting’s midday (ZT4) and midnight (ZT16) serum samples. A total of 112 probable molecules were identified at least in 50% of two groups. PCA analysis using the metadata (112 × 14) including all the probable molecules showed limited overlapping clusters of the two groups (Fig 2C). In a paired-t-test, a set of 12 molecules showed significant variation at p<0.05. Random forest (RF) analysis identified 11 molecules with mean decrease accuracy of more than 0.002. The common set of molecules between t-test, and RF method were identified as important molecules (Fig 2D). Hierarchical clustering also showed distinct separation between these two times of sampling (Fig 2E).

**Figure 2:**
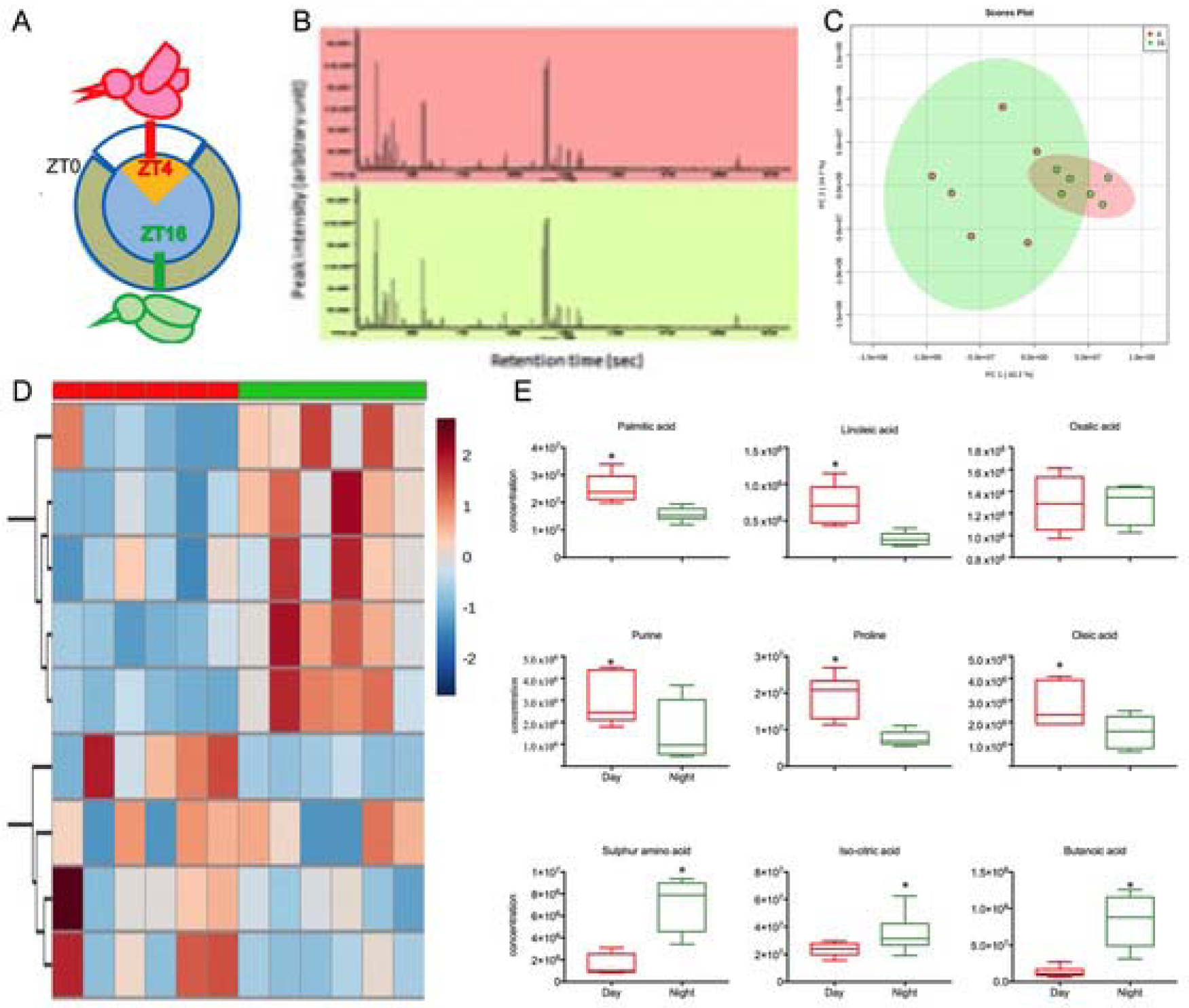
A. Time of blood sampling of male redheaded bunting*, Emberiza bruniceps* held under 8L: 16D. (ZT4, red) and night (ZT16, green). B. Representative total ion chromatograms of derivatized metabolites isolated from day (ZT4, red) and night (ZT16, green) groups. C. Score plot generated from principal component analysis (PCA) of entire data matrix using all the variable shows distinct pattern of day (ZT4, red). and night (ZT16, green) groups. D. Heatmap showing hierarchical clustering of observational and selected 9 variables show distinct cluster of day (ZT4, red). and night (ZT16, green) groups. E. Day (red) and night (green) levels of palmitic acid, linoleic acid, oleic acid, purine, proline, oxalic acid, sulphur amino acid, iso-citric acid and butanoic acid. Asterisk (*) on a bar in (F.1-F.9) indicates a significant difference between midday and mid-night values (p < 0.05, Student t test).

### Serum metabolomic variations between day and night

There was a significant difference in concentrations of between 9 metabolites discriminating day from night. Figure 3 shows significantly higher levels of sixteen and eighteen carbon fatty acids viz. palmitic acid (palmitate, 16:0; p=0.0011), oleic acid (oleate) (18:1; p=0.0325), linoleic acid (linoleate, 18:2, p=0.0022; Mann Whitney test) and linolenic acid (linolenate) (18:3) in the daytime. Purine exhibited day time high in most samples in group, but the difference was not statistically significant. But, amino acid proline significantly (p=0.0011) reduced during the night. Taurine, a sulphur amino acid, implicated in scavenging of reactive carbonyls, was higher in the night (t_5_=4.05; p<0.005)

Higher abundance of Iso-citric acid, was observed in night samples. Few short chain fatty acids, such as, butanoic acid (p=0.0022) and iso-citric acid (p=0.0465), a TCA intermediate, were present in significantly higher amount in night samples. Oxalic acid (p= 0.4429) abundance was higher in the night samples but was statistically insignificant.

### Supplementary results

Within group variation of midday and midnight were found to be similar (p=0.28, Fig S2). Identities of five of these important molecules (oleic acid, palmitic acid, linoleic acid, proline and uric acid) were established by matching retention time and fragmentation pattern of commercial standards (Fig S3). PCA was carried out with a revised matrix with 6 important molecules to verify the clustering pattern. Although similar pattern of PCA score plot was observed with two distinct groups, the new model explained ~84% of variance in the first two components as compared to ~28% variance in the earlier PCA (Fig S4).

## DISCUSSION

Present study compared day and night serum metabolome of wintering redheaded buntings. To date, metabolomics analyses depicting functional association of metabolism with circadian rhythms largely restricted to model organisms like drosophila, mice and humans. Only a single report pertains to day-night changes in fish liver metabolites medaka, *Oryzias latipes* (Fujisawa et al., 2016). Birds, unlike mammals and fish, exhibit discrete daily food eating patterns thus offering added advantage while studying circadian correlates of metabolism (Guglielmo, 2010).

Our behavioural assay of activity and feeding corroborated an earlier proposition that a circadian timing system mediates temporal synchrony of behaviour and physiology (Singh et al., 2015). Overall, buntings followed standard patterns of day-night differences in daily pattern of feeding, activity and body temperature for birds (Morton, 1967). The dawn and dusk peak in food intake may be explained by exponential model as proposed by Polo & Bautista, 2006. In small birds a decrease in fat and water reserves during the night accrues a need for recovery, resulting into an increase in intake at the start of the next foraging period showing the greatest rates of body mass gain at dawn and before dusk. This decreases risk of starvation during the morning when fat reserves are lowest, but putting on weight too might enhance risk of predation; a second peak in foraging activity at dusk permits birds to put on enough fat for the next nocturnal period while minimizing the body mass dependent costs during the day. Active food intake and hence energy utilization during daytime might also be related to the cyclic change in body temperature corresponding to body’s thermogenic demands, and hence the daily metabolic variation (Swanson, 2010).

Our standardized method of serum metabolite profiling, adopted for the first time in buntings aimed to explore well-defined metabolites including those not characterized earlier. Logistically, the present study was restricted to only low energy budgeting stage i.e. premigratory LHS. This could, in future, be useful to monitor the physiological changes in birds during infection, migration, and might be extrapolated to human benefit. In this serum metabolite profiling study, a total of 112 analytes were identified and majority of these molecules belonged to amino acids, monosaccharide, organic and fatty acids. Despite significant diurnal differences in feeding and activity, overall metabolites maintained a homeostatic concentration. Using both uni and multivariate analyses, we identified a set of 6 molecules (palmitic acid, oleic acid, linolenic acid, uric acid, proline and analyte 147@528) that were high in the daytime groups. Higher abundance of these molecules at daytime reflects food intake status and their usefulness to contribute the energy demand.

The narrow range of fatty acid composition observed, was typical for wild passerines and their abundance at a given time was affected in part by diet and selective mobilization depending on state of migration (Egeler & Williams, 2007; Price et al., 2008). In buntings, sixteen and eighteen carbon fatty acids viz. palmitic acid (palmitate) (16:0), oleic acid (oleate) (18:1), linoleic acid (linoleate) (18:2) and linolenic acid (linolenate) (18:3) exhibited daytime elevation. Relative higher abundance of oleic and linolenic acids in midday samples in buntings is in agreement with observations on wintering sparrows; the latter are reported to selectively prefer mobilization of palmitate and linoleate (Price et al., 2008). Total fasting in the night seemed to have triggered lipolysis, as apparent from increased concentrations short chain fatty acid, butanoic acid, detected at night.

Significantly higher levels of iso-citrate, a TCA cycle intermediates were detected in the night. It is possible that TCA cycle was interrupted at this key step because 1) lower body temperature (Fig1H) in the night confounded NAD+/Isocitrate dehydrogenase function, a regulatory enzyme within the tricarboxylic acid cycle, 2) higher levels of aconitase in night (Reddy et al., 2006) converted citric acid to iso-citric acid. More so, fluctuations tending towards higher titres of oxalic acid in the night are suggestive of oxidation of glucose in night fasted birds. This may also be linked to lack of any diurnal variation in glucose abundance in short-day buntings. This absence of circadian rhythm in serum glucose was intriguing in the present study, although indirectly in agreement with earlier studies. For example, presence of enlarged heart concurrent with daytime elevated mRNA levels of malate dehydrogenase (mdh) in premigratory buntings have been reported (Trivedi et al., 2015), which in turn expedites glucose utilization (Razeghi et al., 2001). So, a daily glucose rhythm could’ve been compromised in wintering buntings, by oxidation/utilization of glucose as stated. This cannot rule out a possibility of diurnal orchestration of glucose in migratory stages of bunting’s annual life (Guglielmo et al., 2017). In Malachite sunbirds daily variation in plasma glucose levels affected the circadian alternation of feeding and fasting (Downs et al., 2010).

Uric acid, a proteolysis waste by-product, has free radicals scavenger cum antioxidant potential and is often taken as marker to characterize the physiological state of the birds (Jenni-Eiermann et al., 2002). Wingfield and others, 2005, observed reduction in activity accompanied lowering of serum uric acid levels in bar-tailed godwits, *Limosa lapponica taymyrensis.* In buntings, diurnal variation in uric acid levels might be coupled to diurnal rhythm of activity.

Significantly higher concentrations of taurine, a reactive carbonyl scavenger amino acid, in avian blood as compared to mammals (Szwergold & Miller, 2014) suggested birds as a potential model to serve as pathology-free model of type 2 diabetes. We observed night time elevation of taurine and higher taurine levels are implicated in prevention of retinal regeneration (García-Ayuso et al., 2018). Taurine, as an inhibitory neurotransmitter has anticonvulsant and antianxiety properties and its metabolic role is to promote insulin release. Therefore, taurine might have a putative role in mediating subtle metabolic advantages to night-migrant songbirds such as weight gain loss consequences or resisting glaucoma despite high glucose levels.

Herein, we have explained the diurnal metabolic variations in redheaded buntings exposed to short days using metabolomics analysis. A standardized method for global avian serum metabolite analysis using GC-MS is established. This molecular profiling method could be useful to carry out metabolic phenotyping of pre and post migratory status in buntings and other species, at a later stage. Present study adds comprehensively to increasing understanding in avian migrants that involves interplay of circadian clock, environmental LD cycle and metabolic phenotype.

## DECLARATION OF INTEREST STATEMENT

The authors declare no competing financial interests.

Ethics Statement-Experiments on birds were performed as per the guidelines of Institutional ethical committee of Chaudhary Charan Singh University, Meerut after receiving appropriate approval from Chief Wildlife Warden Rajasthan, Jaipur to work on buntings.

Financial support to NJG from Science and Engineering Research Board, New Delhi, India (SR/SO/AS-69/2011) to conduct experiment and Core fund of International Centre for Genetic Engineering and Biotechnology New Delhi to RKN to conduct experiment, is acknowledged.

