## Supplemental Files for "Diurnal variations in serum metabolites of wintering redheaded buntings"

Supplementary Figure S1:

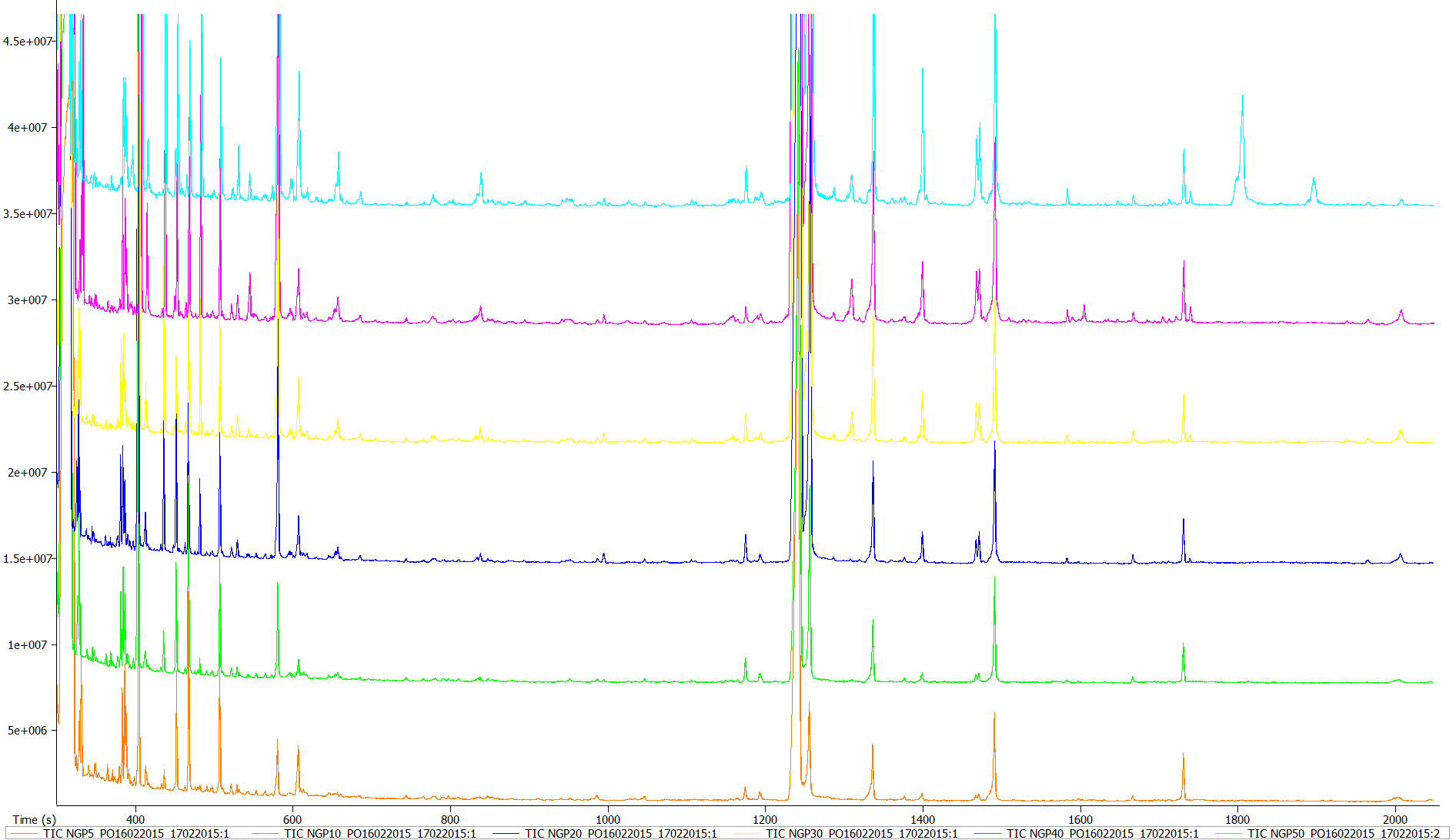

F

E

D

C

B

A

Supplementary Figure S2:

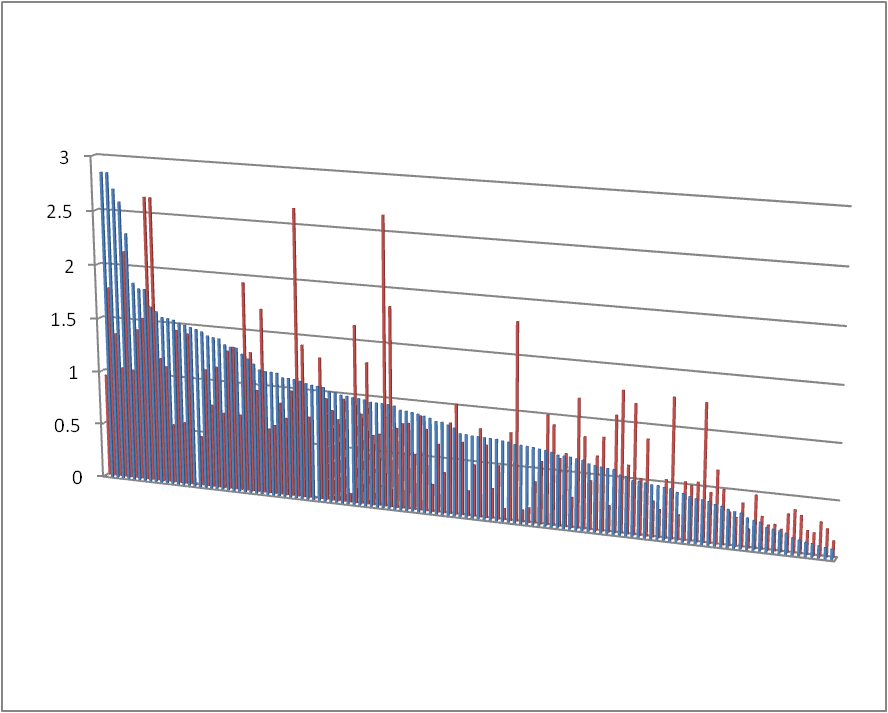

Coefficient of variation

Metabolites

Supplementary Figure S3:

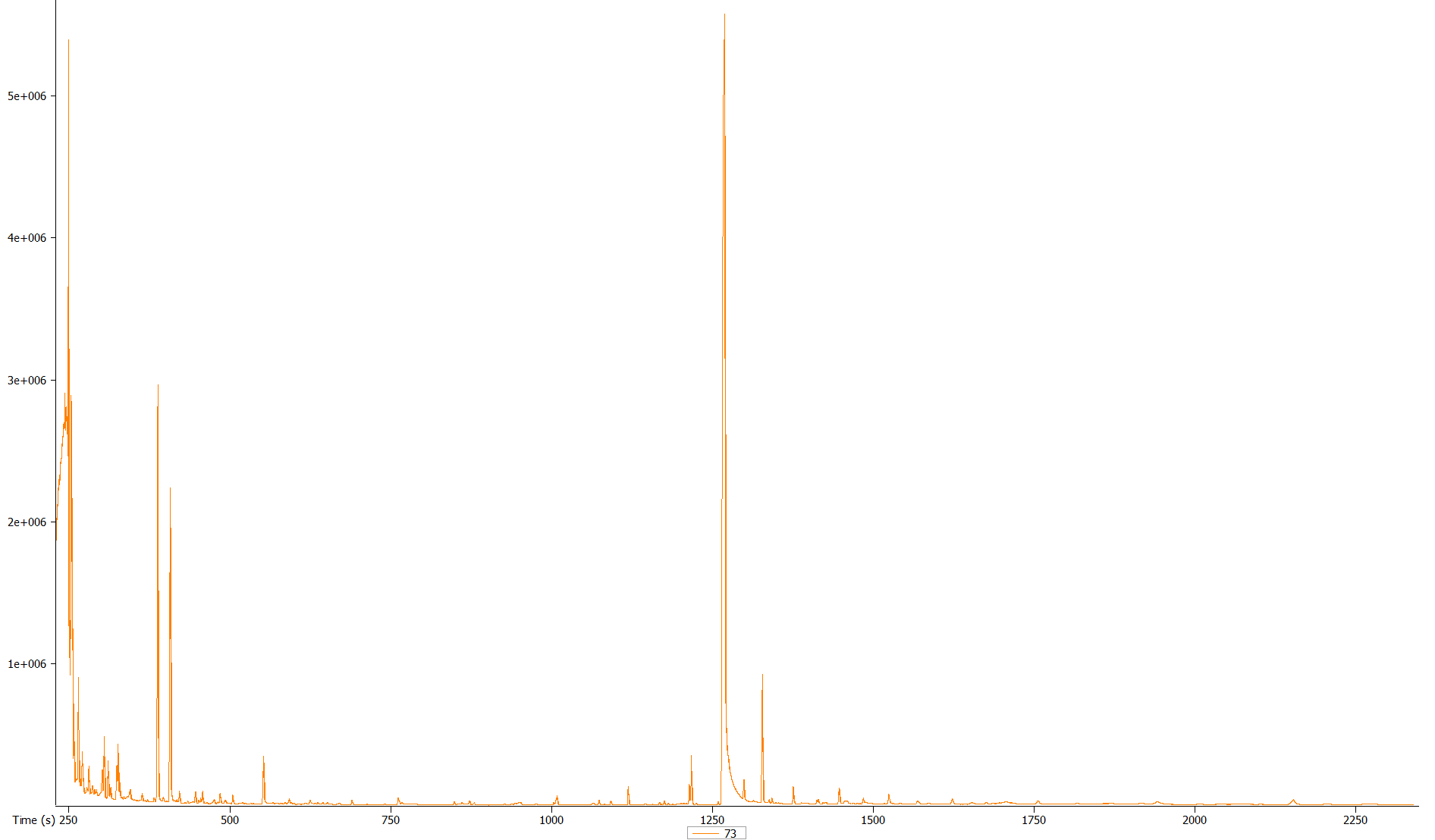

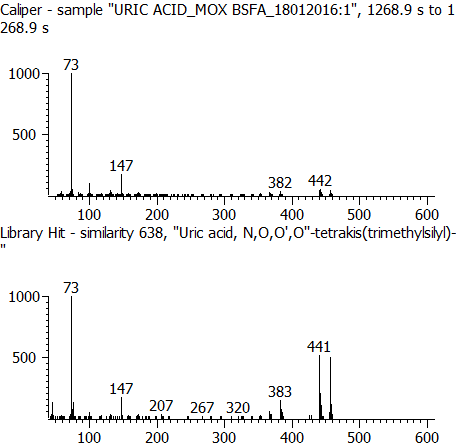

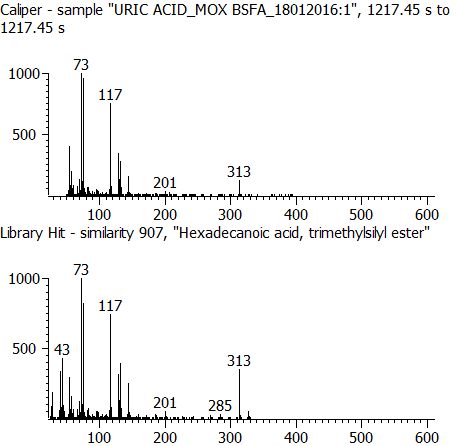

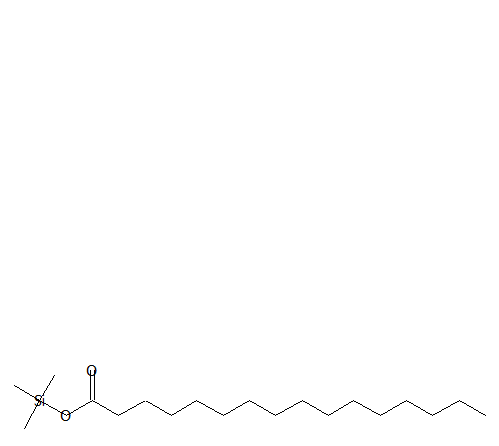

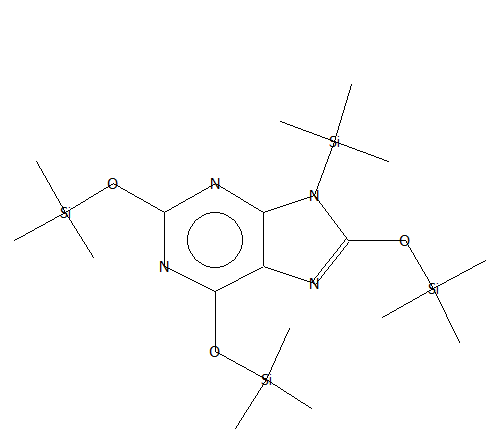

**(A) Hexadecanoic (Palmitic) acid and (B) Uric acid**

**Hexadecanoic (Palmitic) acid**

**Uric acid**

Supplementary Figure S3: continued..

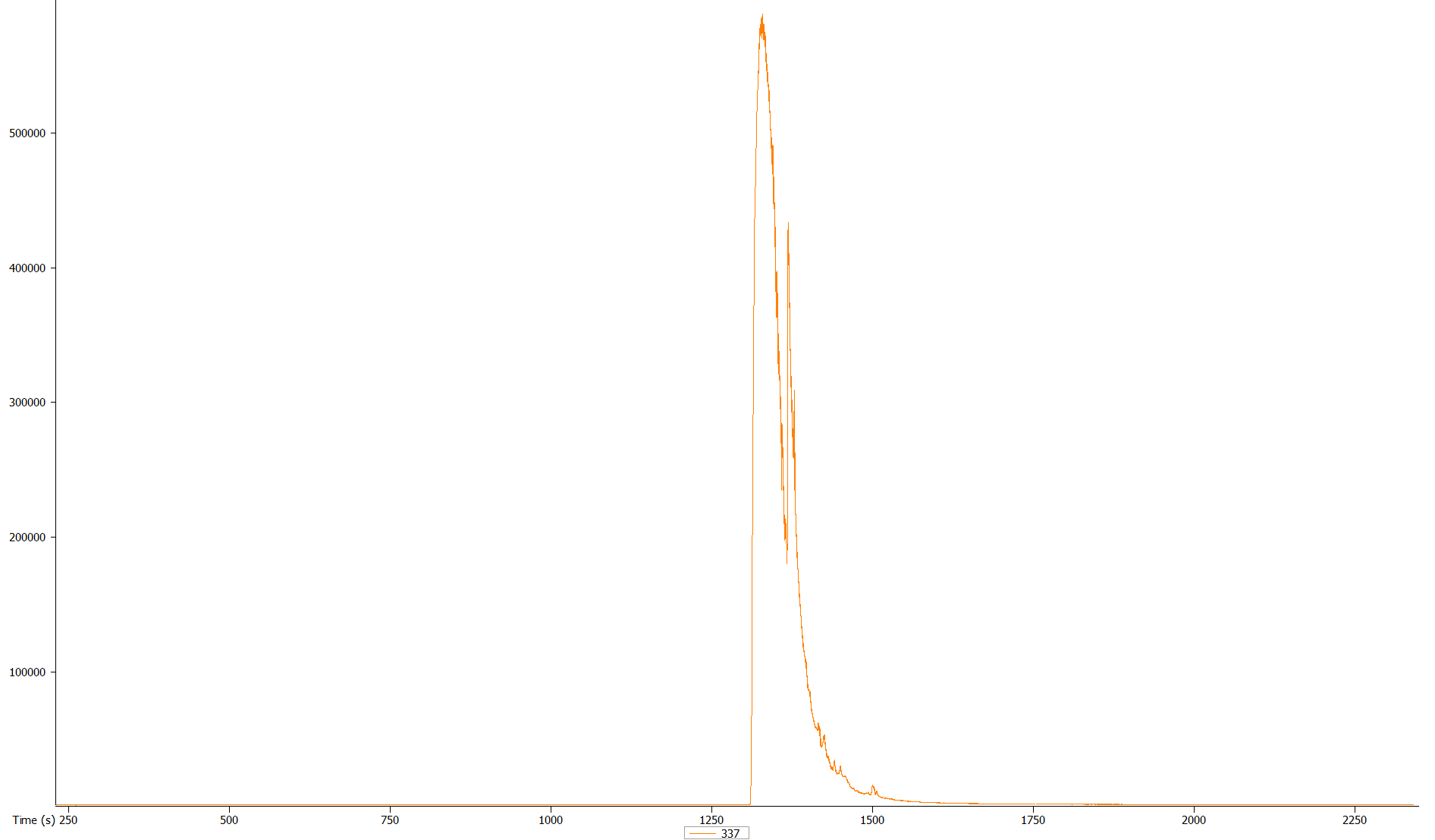

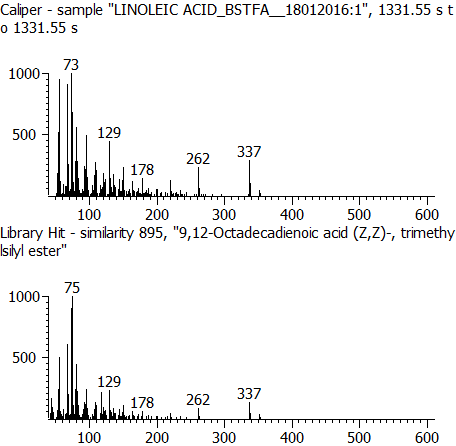

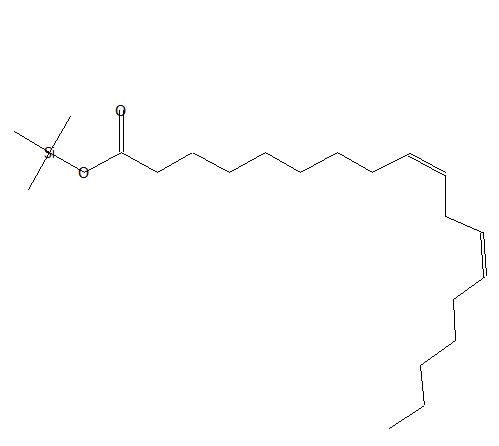

**(C) Linoleic acid**

**Linoleic acid**

Supplementary Figure S3: continued..

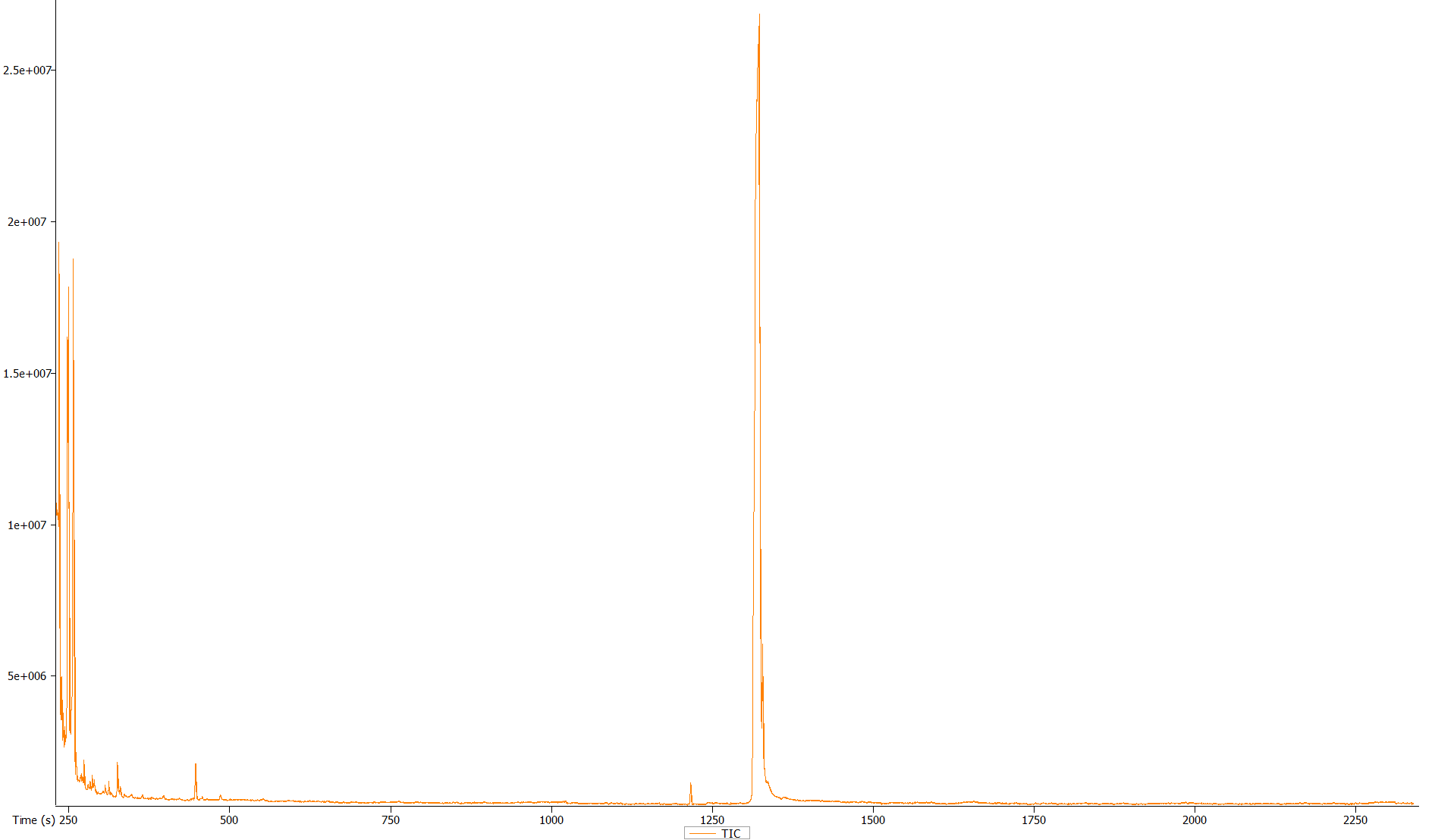

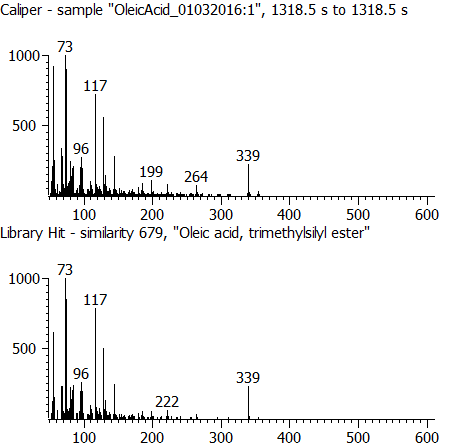

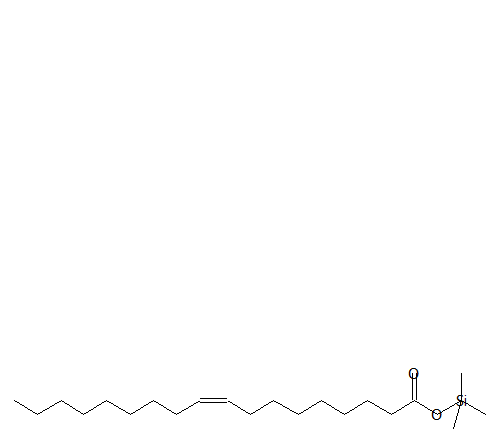

**(D) Oleic acid**

**Oleic acid**

Supplementary Figure S3: continued..

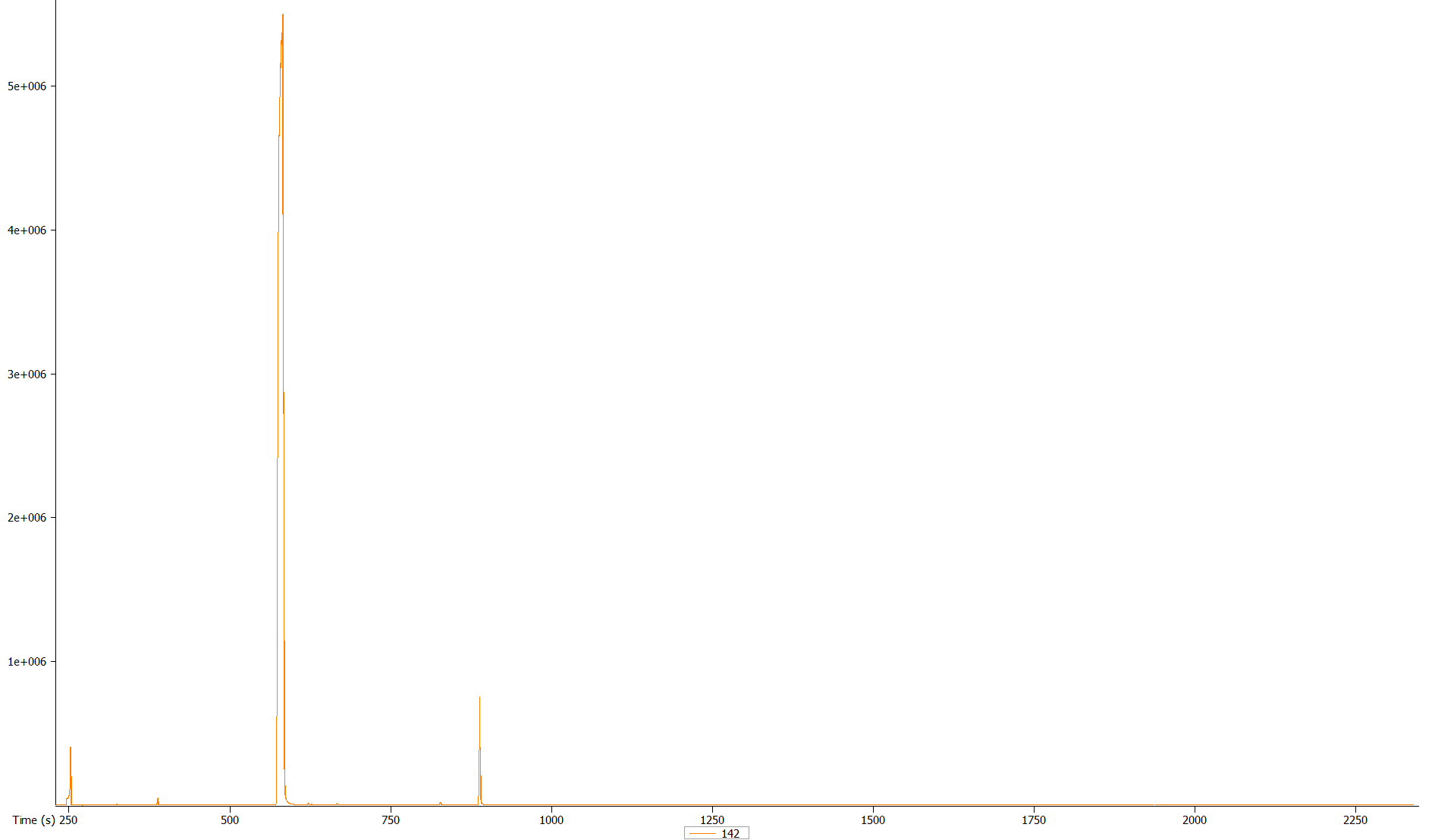

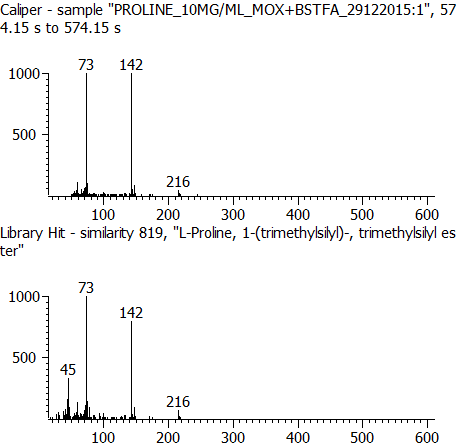

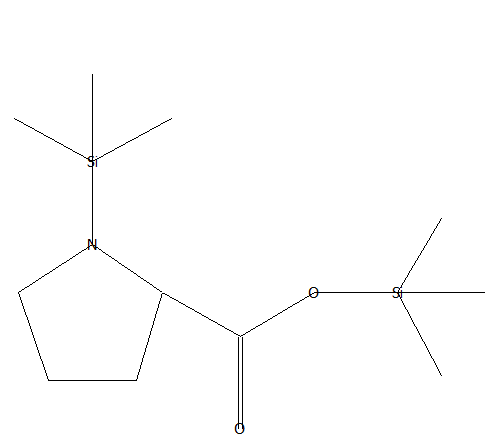

**(E) Proline**

**Proline**

Supplementary Figure S4:

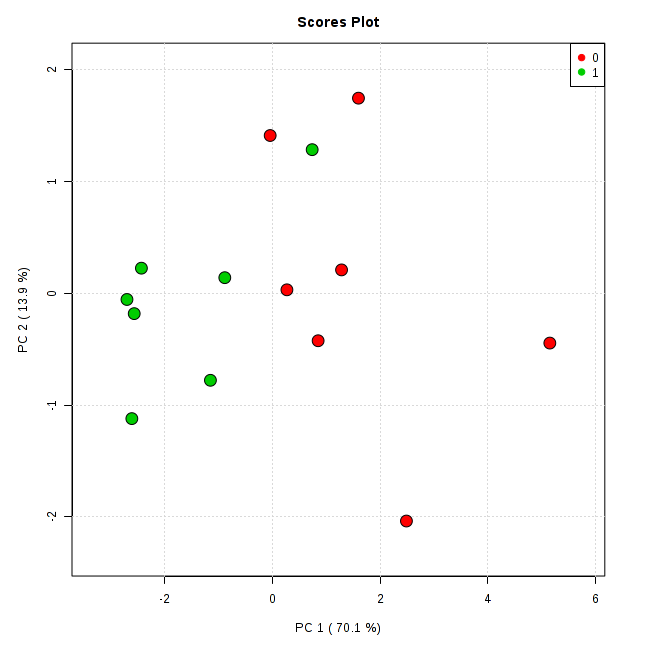

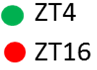

Supplementary Figure S5:

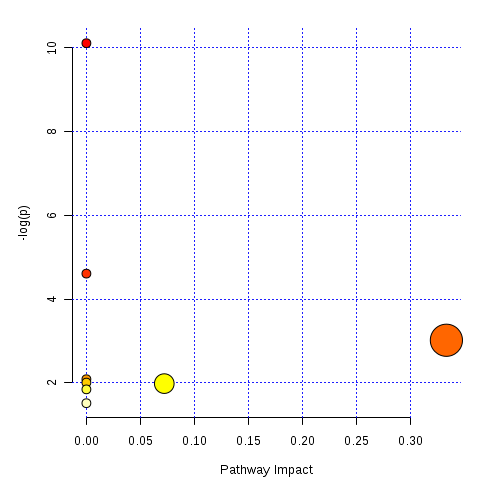

Supplementary Table S1: Common molecules identified in bird serum after GC-MS analysis. Reported retention index (RI) information are from NIST otherwise from Golm database as mentioned.

| **Common name** | **RI Calculated** | **RI Reported** | **Column used in Reported RI** |
| --- | --- | --- | --- |
| 2-Aminobutyric acid | 1182 | 1161.5 | VF-5MS |
| 2-Hydroxyisovaleric acid | 1178 | 1174 | HP-5 |
| 2-Oxoglutaric acid | 1593 | 1587 | 5 % Ph -MS |
| 3-Hydroxybutyric acid | 1169.5 | 1147.9 | VF-5MS |
| 3-Hydroxyisovaleric acid | 1219 | 1214 | DB-5 |
| Alpha-ketoisovaleric acid | 1118 | 1096 | VAR5 (Golm) |
| Aminomalonic acid | 1492 | 1485 | Methylsiloxane, 5 % Ph groups |
| Arachidonic acid | 2380 | 2393 | HP-5 |
| Aspertic acid | 1543 | 1534 | HP-5MS |
| Benzoic acid | 1254 | 1253 | DB-5 |
| Butan | 1322 | 1314 | DB-5 |
| Citric acid | 1845.9 | 1838 | DB-5 |
| Docosahexaenoic acid (DHA) | 2568 | 2562 | DHA-TMS VF-5MS |
| Ethanolamine | 1042 | 1027 | HP-5MS |
| Ethylene glycol | 1006 | 990 | HP-5 |
| Galactose | 1911 | 1897.6 | VF-5MS |
| Glucose | 1969 | 1953 | OV-1 |
| Glutamic acid | 1638 | 1629 | OV-1 |
| Glycerol | 1289 | 1282 | DB-5 |
| Glycine | 1127 | 1105 | DB-5 |
| Glycolic acid | 1083 | 1078 | DB-5 |
| Hypoxanthine | 1820 | 1812.2 | VF-5MS |
| Isoleucine | 1306 | 1300 | DB-5 |
| Leucine | 1284 | 1276 | DB-5 |
| Linoleic acid | 2218 | 2212 | HP-5 |
| Malic acid | 1512 | 1480 | VAR5 (Golm) |
| Mannonic acid | 2015 | 2012 | OV-1 |
| Myristic acid | 1853 | 1850 | HP-5 |
| Oleic acid | 2224 | 2224 | HP-5 |
| Ornithine | 1632 | 1624 | DB-5 |
| Oxalic acid | 1150 | 1131 | DB-5 |
| Oxo-Proline | 1548 | 1524.2 | DB-5MS |
| Palmitelaidic acid | 2032 | 2030 | HP-5MS |
| Palmitic acid | 2052 | 2050 | HP-5 |
| Phenylalanine | 1643 | 1640 | DB-5 |
| Proline | 1541 | 1524.2 | DB-5MS |
| Ribitol | 1759 | 1766 | HP-5 |
| Sarcosine | 1148 | 1143 | OV-1 |
| Serine | 1375 | 1380 | OV-1 |
| Stearic acid | 2249 | 2239 | HP-5 |
| Succinic acid | 1345 | 1336 | DB-5 |
| Taurine | 1690 | 1676 | HP-1 |
| Threonic acid | 1586 | 1545 | VAR5 (Golm) |
| Threonine | 1402.4 | 1367.4 | VF-5MS |
| Tyrosine | 1963 | 1958 | DB-5 |
| Uric acid | 2135 | 2127 | DB-5 |
| Valine | 1127 | 1221 | DB-5 |
